# Evaluation of phage activity against diverse *Enterococcus faecalis* from infective endocarditis

**DOI:** 10.64898/2026.07.24.740664

**Authors:** Yanhong Li, Madison J. Moreland, Kirsten M. Evans, Madison E. Stellfox, Daria Van Tyne

**Affiliations:** Division of Infectious Diseases, University of Pittsburgh School of Medicine, Pittsburgh, PA; School of Clinical Medicine, Tsinghua Medicine, Tsinghua University, Beijing, China

**Keywords:** *Enterococcus faecalis*, phage therapy, infective endocarditis

## Abstract

*Enterococcus faecalis* is a leading cause of infective endocarditis, particularly in older and hospitalized patients. Treatment is challenging due to the intrinsic resistance of *E. faecalis* to many clinically important antibiotics and the increasing prevalence of multidrug-resistant strains. Bacteriophage (phage)-based therapies, including phage cocktails and phage-antibiotic combinations, offer a promising approach for treating resistant bacterial infections. In this study, we isolated ten novel phages from wastewater by screening on *E. faecalis* clinical isolates and tested phage activity both individually and in phage-phage and phage-antibiotic combinations against 18 *E. faecalis* clinical isolates collected from patients with infective endocarditis. We found that 15/18 (83%) isolates were susceptible to at least one phage tested, and that the same proportion of isolates were susceptible to a cocktail of three phages with complementary activities. We also observed improved bacterial growth inhibition and increased killing when phages were combined with ampicillin or daptomycin, two antibiotics commonly used to treat *E*. *faecalis* infections. Finally, we found that treatment with the three-phage cocktail improved the survival of *Galleria mellonella* infected with a pathogenic *E. faecalis* endocarditis isolate. Overall, these findings suggest that phages could be a helpful addition to the therapeutic repertoire for treating *E. faecalis* infective endocarditis.

**Importance:** New approaches are needed for treating infective endocarditis, a life-threatening heart infection. Here we focus on *Enterococcus faecalis*, a resilient bacterium that is increasingly resistant to standard-of care antibiotic regimens, and explore the use of phage therapy for treating *E. faecalis* endocarditis. We found that combining phages together in a cocktail or pairing them with existing antibiotics can effectively kill *E. faecalis* clinical isolates from patients with infective endocarditis, and that treatment of infected waxworm moth larvae with a phage cocktail improved their survival. The results of this study contribute to the development of phage-based therapeutic approaches for *E. faecalis* endocarditis, which could ultimately lead to improved outcomes for patients facing this severe infection.

## Introduction

Infective endocarditis (IE) is a serious infection of the heart valves and endocardium that is typically caused by bacterial pathogens. Despite advances in diagnosis and treatment, IE mortality remains alarmingly high, reaching up to 24%^1^. Moreover, the incidence of IE is increasing, exceeding expected trends^2^. Enterococci, a genus of Gram-positive commensals that live in the gastrointestinal tract, are a frequent cause of IE, especially in elderly and immunocompromised patients^1^. Enterococcal IE accounts for 13% of IE cases, and has been associated with a higher relapse rate compared to IE caused by other pathogens^3^.

Of the dozens of known enterococcal species, *Enterococcus faecalis* is responsible for 90% of enterococcal IE cases, and poses a significant treatment challenge^4^. *E. faecalis* IE (EFIE) often results from healthcare exposure, genitourinary procedures, and long-standing colonization^5^. *E. faecalis* strains possess both intrinsic and acquired resistance to many clinically important antibiotics, like cephalosporins and aminoglycosides, and continue to develop multidrug-resistance^6^. In addition, *E. faecalis* can form biofilms, which are communities of microbes that stick to abiotic and biotic surfaces using a polymeric matrix, thereby decreasing the ability of antimicrobials to penetrate and kill bacterial cells^7^. Single-agent antibiotic therapies often fail to effectively treat EFIE, and multi-drug regimens are the standard of care^4^. Current guidelines recommend ampicillin plus gentamicin (AG) or ampicillin plus ceftriaxone (AC) as treatment for EFIE, with AC being preferred due to its comparable efficacy to AG but better renal safety and fewer adverse events^8^. However, the mortality of EFIE still remains as high as 30%^4^, and recent clinical isolates from patients with EFIE have demonstrated reduced susceptibility to both ceftriaxone alone and the AC combination^9,10^. Daptomycin, a lipopeptide that disrupts cell membranes, has been used in some EFIE cases when AC is not adequately effective or in patients who cannot tolerate AC^11^. However, daptomycin resistance was reported in 5.1% of *E. faecalis* isolates in a recent study^12^. These findings raise concerns about the efficacy of current treatment regimens and highlight the need for alternative therapeutic strategies for EFIE.

Bacteriophages, or phages, are viruses that specifically infect and kill bacterial cells^13^. The host range of a phage, or the spectrum of bacterial species or strains it can infect, can vary widely. Some phages exhibit a broad host range, capable of infecting multiple related species, while others are highly specific, targeting only a single bacterial strain. Due to their ability to effectively lyse bacterial cells, lytic phages have been explored as potential adjuvants and alternatives to antibiotics for the treatment of antibiotic-resistant infections^14^. Prior studies have identified several phages that demonstrate effective lytic activity against *E. faecalis*, in both planktonic and biofilm conditions, and regardless of bacterial antibiotic resistance profiles^15^. However, as with antibiotics, bacteria can develop resistance to phages through multiple strategies such as mutations in the phage receptor, receptor masking, and anti-phage defense genes^16^. As a potential way to address this, phage cocktails containing combinations of two or more phages have been developed to broaden phage host range and mitigate resistance by targeting multiple bacterial receptors^15,17^. Phage cocktails have been shown previously to successfully eradicate multidrug-resistant *E. faecalis* in both *in vitro* and *in vivo* models^18,19^. In addition, combining phages with antibiotics can produce synergistic effects that enhance bacterial clearance *in vitro*^15,20,21^. These prior studies have generated interest among researchers and clinicians regarding the use of phage therapy for *E. faecalis* infections. However, the potential of utilizing phage therapy in treating EFIE has not been fully explored.

In this study, we isolated and characterized 10 new *E. faecalis*-targeting phages from municipal wastewater and evaluated their activity against 18 EFIE isolates both *in vitro* and in a *Galleria mellonella* (waxworm) model of *E. faecalis* infection. We observed distinct phage susceptibility patterns among diverse EFIE isolates, found that phage-ampicillin or phage-daptomycin combinations generally outperformed monotherapy, and designed a three-phage cocktail that exhibited robust and broad-spectrum activity. Overall, this study shows the therapeutic potential of phage-based approaches to address antibiotic-resistant *E. faecalis* infections such as EFIE.

## Results

### Genomic characterization of 10 new *E. faecalis-*targeting phages

We isolated 10 *E. faecalis*-targeting phages using four different *E. faecalis* clinical isolates to screen for lytic phages from municipal wastewater collected in the Pittsburgh area (**Table 1**). Most phages had genomes of 40-42 kb, except EFS15, which had a significantly larger genome of nearly 149 kb. GC content was similar across the 9 phages with smaller genomes (34.3-35.0%), while EFS15 had a higher GC content of 37.3% (**Table 1**). Taxonomic analysis revealed that 5/10 (50%) phages were predicted to belong to the *Efquatrovirus* genus, which has been previously described to include phages that target *E. faecalis*^22,23^. The remaining phages belonged to unknown genera, and their placement on a phage genome phylogeny suggested that the isolated phages likely belong to four different genera (**Fig. 1**). All 10 phages were predicted to have virulent lifestyles^24^ (**Table 1**), indicating their potential suitability for therapeutic applications. We also investigated the species-specificity of the isolated phages by testing their activity against six previously characterized *E. faecium* clinical isolates (**Table S1**). We did not observe phage activity on any *E. faecium* isolates, suggesting that the phages we isolated are only able to infect *E. faecalis*.

**Figure 1.**
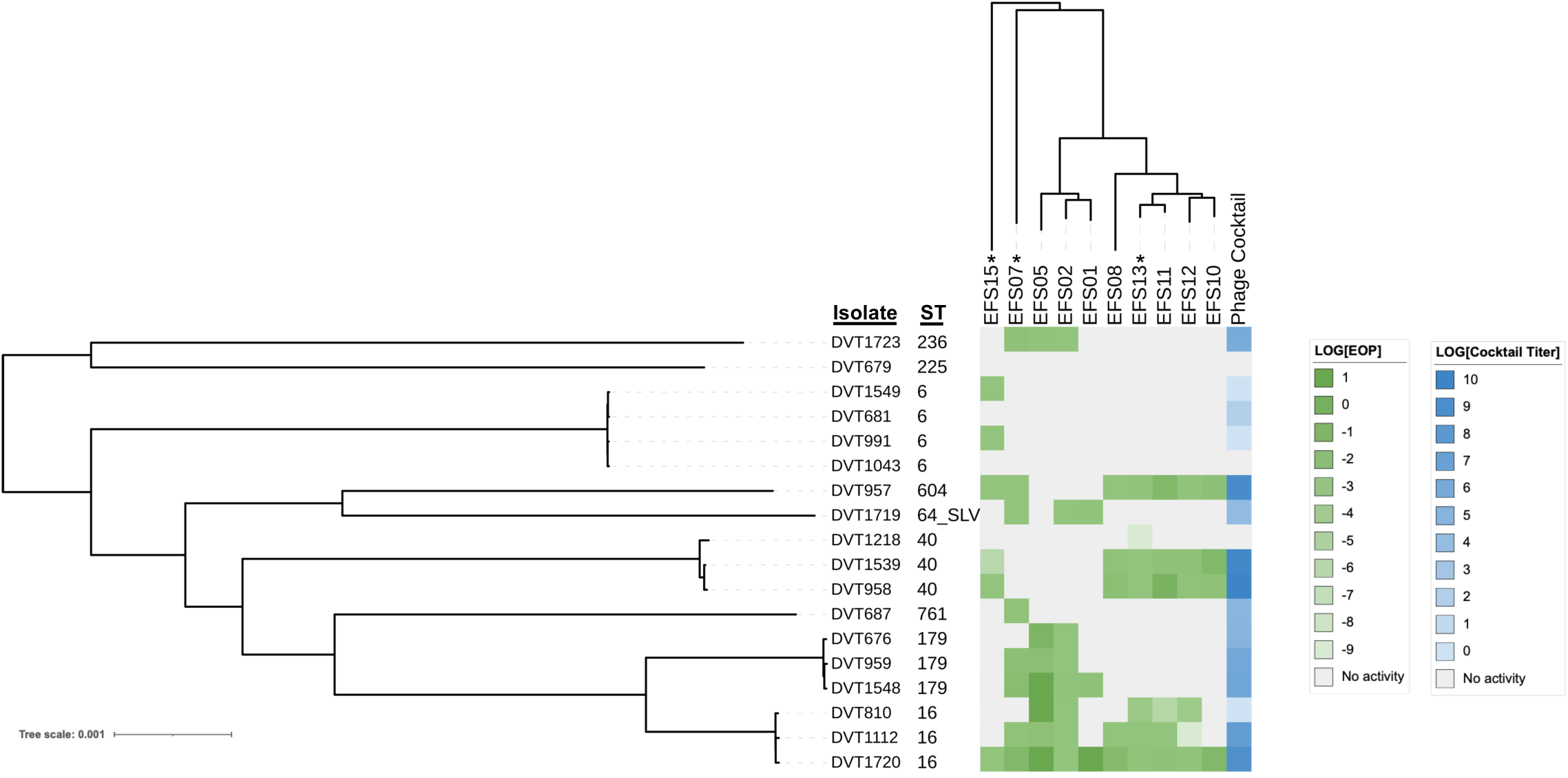
Bacterial and phage genomic diversity and and phage host range across clinical EFIE isolates. A midpoint-rooted, core genome phylogenetic tree of 18 EFIE isolates was constructed with RAxML. Tips are labeled with isolate name and multi-locus sequence type (ST). SLV = single locus variant. The phage dendrogram was constructed with ViPTree using assembled phage genomes. Three phages selected for inclusion in a cocktail are marked with asterisks. Heatmaps display the phage susceptibility of each EFIE isolate. Individual phages and a three-phage cocktail (EFS07+EFS13+EFS15) were tested against each isolate. Green = efficiency of plating (EOP) values; blue = phage cocktail titer; gray = no detectable phage activity. For all colors, darker shades indicate higher values.

**Table 1.**
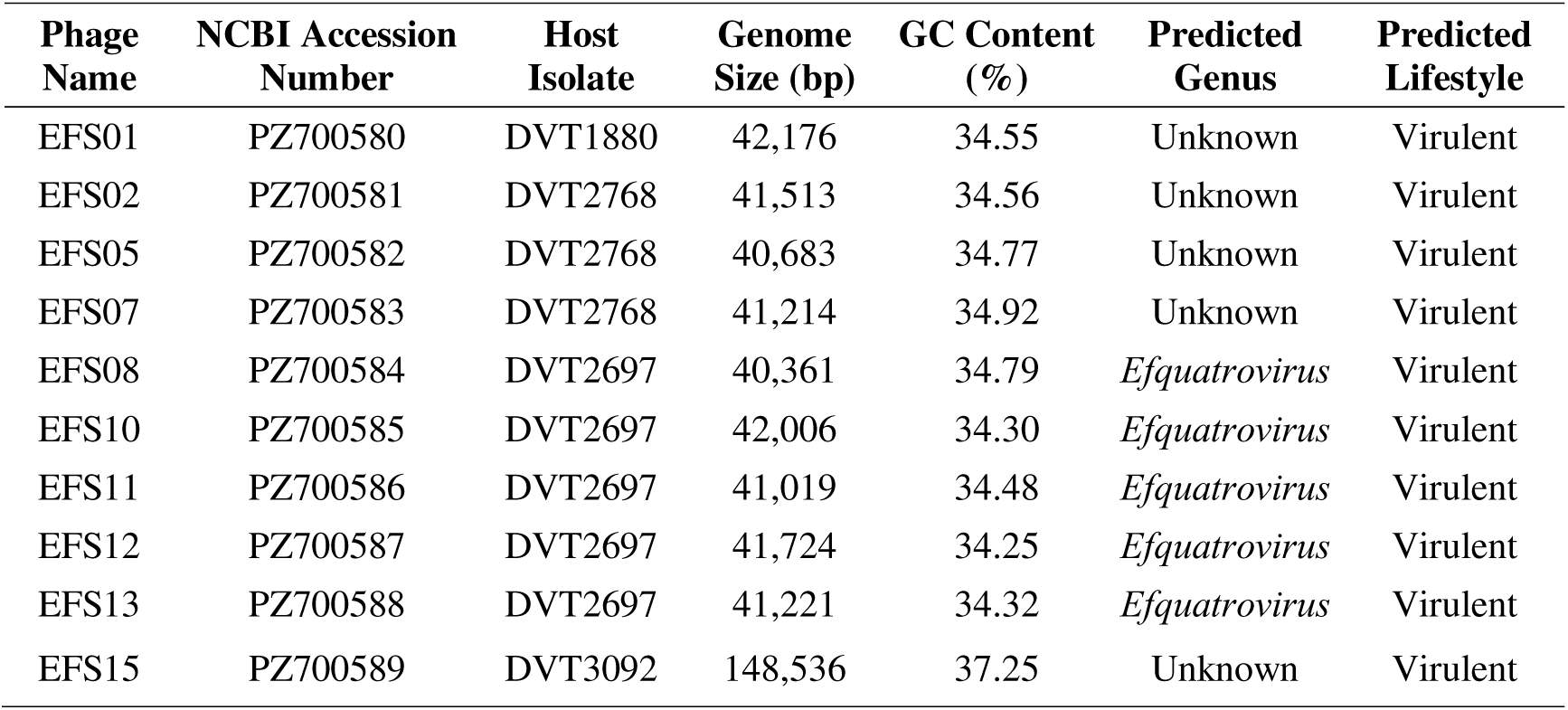
Genome diversity of *Enterococcus faecalis*-targeting phages.

### Diverse phage susceptibility patterns among clinical EFIE isolates

To better understand phage-host interactions across diverse clinical *E. faecalis* isolates, we selected 18 EFIE isolates representing nine different multi-locus sequence types (STs) and evaluated their susceptibility to all 10 *E. faecalis*-targeting phages (**Fig. 1**, **Table S1**)^10^. There was considerable variability in the efficiency of plating (EOP) across different isolate-phage combinations, however several isolates showed broad phage susceptibility. Isolates in the ST16 lineage were particularly phage-susceptible, with one ST16 isolate (DVT1720) displaying sensitivity to all phages tested. ST40 and ST179 isolates exhibited variable phage susceptibility, while ST6 isolates appeared to be the least phage-susceptible. Each individual phage was only active against a subset of isolates, with individual phage activities ranging from 3-8 isolates. Next, we selected three phages with broad and complementary activities (EFS07, EFS13 and EFS15), combined them in equal proportion to generate a three-phage cocktail, and then tested the cocktail’s activity against all 18 isolates. The cocktail demonstrated activity against 15/18 (83%) isolates, and susceptibility to the cocktail was similar to individual phage susceptibility. These results suggest that combining different phages into a cocktail increased the spectrum of phage activity, and that the phages in the cocktail did not display antagonism with one another (**Fig. 1, Table S1**).

### Phage-antibiotic combination therapies demonstrate enhanced *in vitro* activity against *E. faecalis*

To evaluate the therapeutic potential of phage-antibiotic combinations against *E. faecalis*, we assessed *in vitro* bacterial growth inhibition over 20 hours using the three phages from the above cocktail and their corresponding host isolates (EFS07 on DVT2768, EFS13 on DVT2697, and EFS15 on DVT3092) both with and without ampicillin and daptomycin, two antibiotics that are frequently used to treat *E. faecalis* infections. We measured bacterial growth dynamics in each condition and used the resulting data to calculate area under the curve (AUC) values for each treatment condition. Most isolates displayed partial growth inhibition with ampicillin or daptomycin alone (tested at concentrations below the minimum inhibitory concentration, MIC) or phage alone (tested at a multiplicity of infection, MOI, of 10), as evidenced by delayed or diminished growth and lower AUC values (**Fig. 2**). In most cases, the combination of antibiotic and phage resulted in significantly less growth than treatment with either agent alone. We also assessed bacterial growth and killing by quantifying colony forming units (CFU) at the beginning and end of each experiment. We observed that ampicillin plus EFS07 or EFS15 resulted in bacterial killing (i.e., CFU reduction) while all other conditions did not (**Fig. S1**). Taken together, these data suggest that phage-antibiotic combination therapies are at least as active as monotherapy with phage or antibiotic alone, and point to positive interactions between these agents.

**Figure 2.**
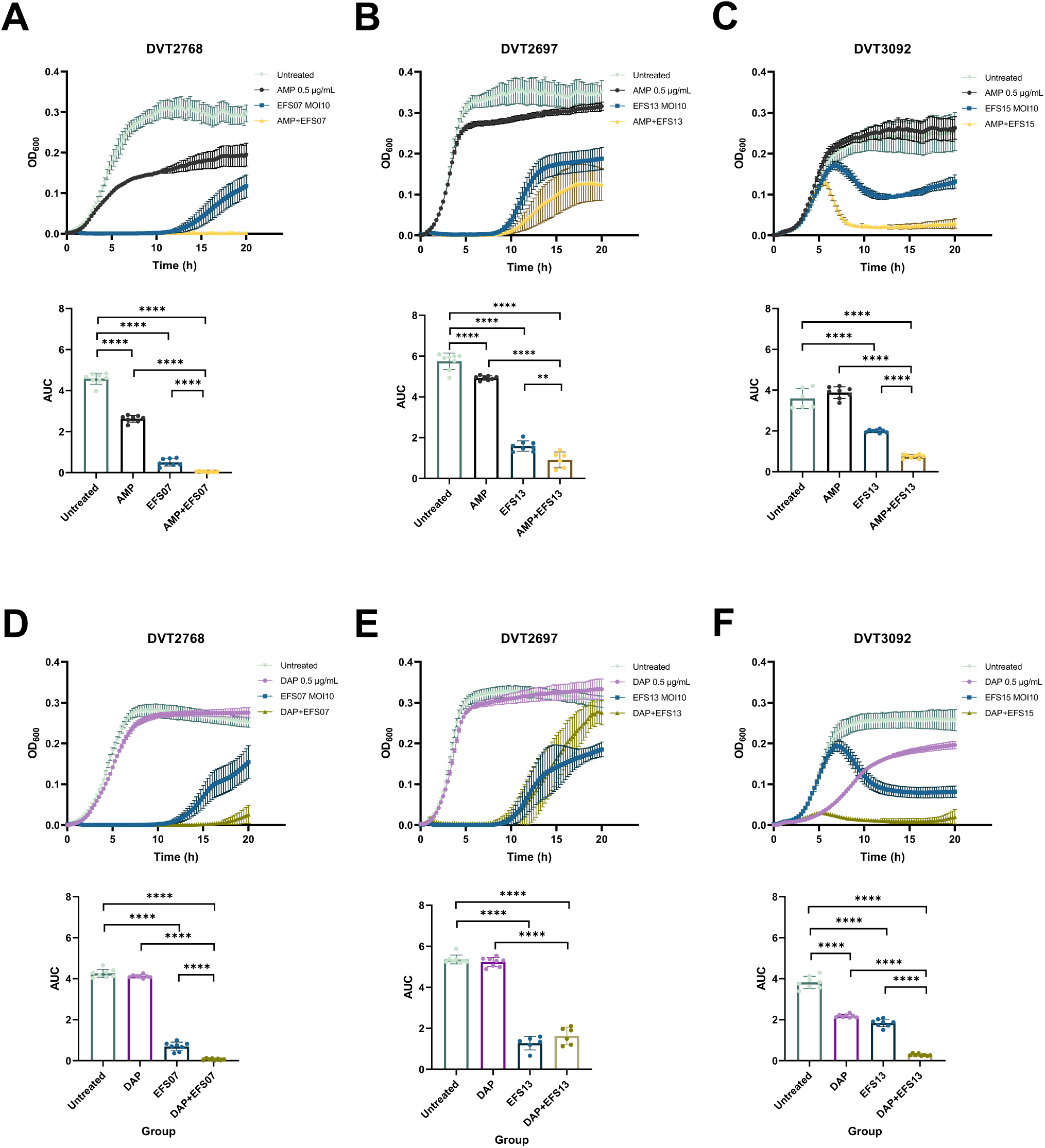
Antibiotic and phage activity alone and together against *E. faecalis* host isolates. Growth curves (upper panels) and area under the curve (AUC) analyses (lower panels) for three *E. faecalis* phage host isolates treated with phages EFS07, EFS13, or EFS15, without or with ampicillin (AMP; panels A-C), or daptomycin (DAP; panels D-F), and grown for 20 hours. DVT2768 was treated with phage EFS07, DVT2697 with phage EFS13, and DVT3092 with phage EFS15 at a multiplicity of infection (MOI) of 10. AUC values represent total bacterial growth over the 20-hour experimental period. Error bars indicate standard error of the mean from six replicates. Statistical significance was determined by one-way ANOVA followed by Tukey’s multiple comparisons test. **p<0.01, ****p<0.0001.

### Phage-antibiotic combination therapy leads to enhanced suppression of EFIE clinical isolate growth

To assess the potential of phage-antibiotic combination therapy against non-phage host EFIE clinical isolates, we monitored the growth dynamics of four isolates representing the four most frequently observed EFIE lineages, including DVT958 (ST40), DVT1548 (ST179), DVT1549 (ST6), and DVT1720 (ST16)^10^. All untreated control conditions showed robust growth throughout the 20-hour incubation period, while ampicillin monotherapy (0.5 μg/mL, 0.5x MIC) modestly reduced bacterial growth and daptomycin monotherapy (0.5 μg/mL, 0.125-0.25x MIC) had no impact, perhaps due to higher daptomycin MICs among these isolates (**Fig. 3A, Table S2**). The three-phage cocktail containing EFS07, EFS13, and EFS15 at an MOI of 100 significantly reduced the growth of most isolates compared with the untreated control condition, however regrowth usually occurred after 10–12 hours, suggesting incomplete clearance and the potential emergence of phage resistance. Notably, in this liquid assay the growth of the ST6 isolate DVT1549 was not inhibited by the cocktail alone, despite the isolate being susceptible to both EFS15 and the cocktail when measured by plaque assay (**Fig. 1**). As expected, combination treatment with the phage cocktail and either ampicillin or daptomycin resulted in near-complete suppression of growth for three isolates (DVT958, DVT1548 and DVT1720). We also observed a modest but significant reduction in the growth of DVT1549 compared with either agent alone, suggesting a positive interaction between the three-phage cocktail and both ampicillin and daptomycin, even though phage susceptibility was minimal in liquid (**Fig. 3B**). Taken together, these findings demonstrate that phage-antibiotic combinations exert improved antimicrobial activity against genetically diverse EFIE clinical isolates, and suggest that this strategy could effectively suppress bacterial growth even against isolates showing reduced susceptibility to individual therapies.

**Figure 3.**
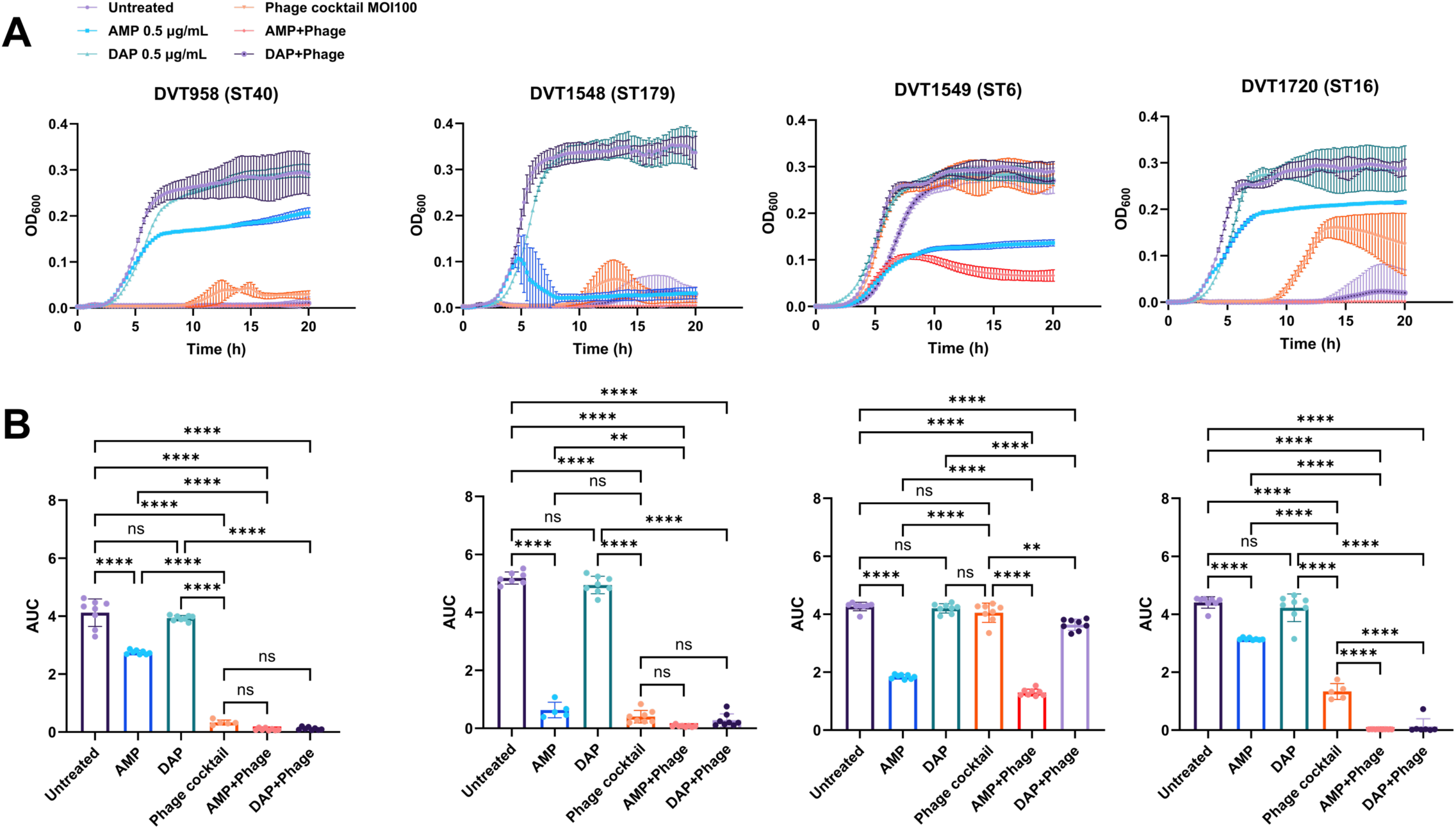
Antibiotic and phage activity against representative EFIE isolates. (A) Growth curves (upper panels) and (B) area under the curve (AUC) analyses (lower panels) for four EFIE isolates treated with ampicillin, daptomycin, three-phage cocktail (EFS07, EFS13, EFS15), or combination therapy over 20 hours. Phage multiplicities of infection (MOIs) of 100 and antibiotic concentrations of 0.5 μg/mL were used in all experiments. AUC values represent total bacterial growth over the 20-hour experimental period. Error bars indicate standard error of the mean of eight replicates. Statistical significance was determined by one-way ANOVA followed by Tukey’s multiple comparisons test. **p<0.01, ****p< 0.0001.

### Three-phage cocktail improves survival and reduces pathogenicity against a multidrug-resistant *E. faecalis* EFIE isolate *in vivo*

To assess the potential *in vivo* efficacy of the three-phage cocktail containing EFS07, EFS13, and EFS15, groups of n=20 *Galleria mellonella* larvae were first infected with 5×10^3 CFUs of each of the four representative EFIE clinical isolates (DVT958, DVT1548, DVT1549, and DVT1720) (**Fig. 4A**). We observed significant differences in three-day survival among the different experimental groups. Compared to the PBS control condition, DVT958 (ST40) and DVT1549 (ST6) caused mortality of ∼50% of worms after three days (DVT958 vs. PBS p<0.01; DVT1549 vs. PBS p<0.0001). In contrast, the DVT1548 (ST179) and DVT1720 (ST16) isolates were less lethal at the same inoculum, with mortality rates between 20 and 30% after three days (**Fig. 4A**). Consistent with increased bacterial virulence, pathogenicity score index (PSI)^25^ values were significantly elevated in the DVT958 and DVT1549-infected groups across all days (**Fig. 4B**). To evaluate the therapeutic efficacy of the three-phage cocktail, we treated worms infected with the DVT1549 isolate with a phage MOI of 100 at two hours post-infection, and compared survival to the PBS-treated and uninfected groups (n=30 worms per group). The DVT1549 isolate was chosen because it exhibited high ceftriaxone resistance and reduced AC combination activity in a prior study^10^, was susceptible to the three-phage cocktail by plaque assay (**Fig. 1**), and caused the highest mortality of *G. mellonella* larvae (**Fig. 4A**). Phage cocktail treatment significantly improved larval survival compared to untreated larvae, with only ∼25% of treated larvae dying by day three compared with over 50% of untreated larvae (p < 0.05, **Fig. 4C**). The phage cocktail also reduced bacterial pathogenicity, with PSI scores among phage-treated larvae significantly lower than untreated larvae on day 1 (p<0.05, **Fig. 4D**). While complete rescue with phage treatment was not achieved, these data nonetheless suggest that the three-phage cocktail can increase survival and reduce bacterial virulence during infection with a multidrug-resistant *E. faecalis* ST6 EFIE isolate.

**Figure 4.**
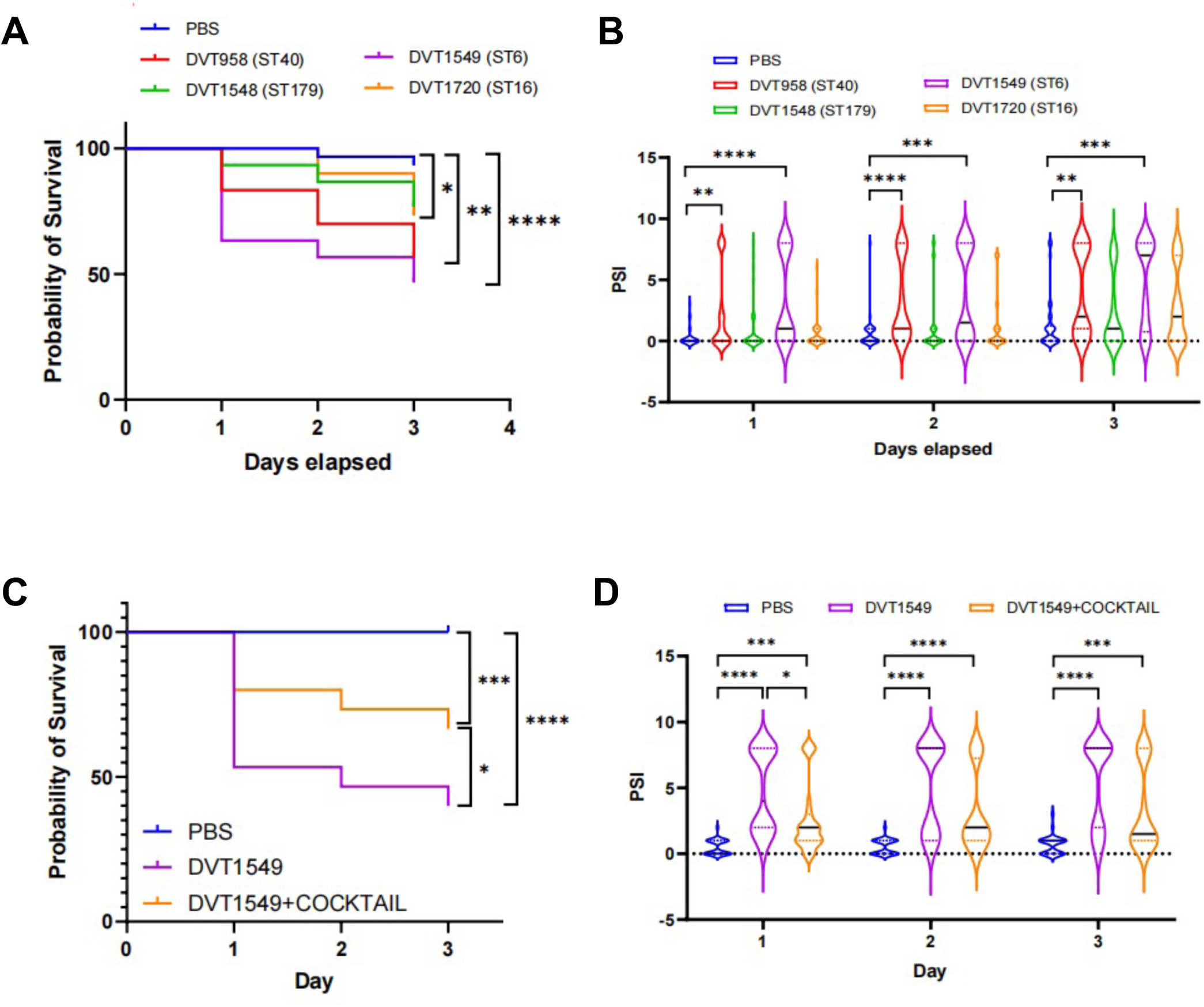
Phage cocktail efficacy *in vivo*. (A) Kaplan-Meier survival curves showing probability of survival over three days for groups of n=20 *Galleria mellonella* larvae injected with 5×10^3^ colony-forming units (CFUs) of four representative EFIE clinical isolates. Statistical significance was determined by log-rank test. (B) Pathogenicity Score Index (PSI) measurements of worms shown in (A) across the three-day experiment. Violin plots show distribution of individual measurements with black lines showing median and dashed lines showing interquartile ranges. Statistical significance was determined by one-way ANOVA followed by Tukey’s multiple comparisons test. (C) Kaplan-Meier survival curves showing probability of survival over three days for groups of n=30 *G. mellonella* larvae injected with PBS, DVT1549 (5×10^3^ CFUs), or DVT1549 followed by phage cocktail (EFS07, EFS13, EFS15) at an MOI of 100 after two hours. (D) PSI measurements of worms shown in (C) across the three-day experiment. Violin plots show distribution of individual measurements with black lines showing median and dashed lines showing quartile ranges. Statistical significance was determined by one-way ANOVA followed by Tukey’s multiple comparisons test. *p<0.05, **p<0.01, ***p<0.001, ****p< 0.0001.

## Discussion

Given the persistently high mortality rates among EFIE patients and increasing antibiotic resistance among clinical *E. faecalis* isolates, novel therapeutic options are needed^4^. The discovery and characterization of new lytic *E. faecalis*-targeting phages is important, as phages can serve as alternative or adjunctive agents to traditional antibiotic therapies. By combining phages with complementary host ranges into a multi-phage cocktail, testing phage activity in combination with antibiotics frequently used to treat *E. faecalis* infections, and validating phage cocktail efficacy in an *in vivo* model of infection, our results suggest that phage-based therapies may constitute an effective strategy that can broaden our antimicrobial armamentarium against *E. faecalis*.

Most previously described *E. faecalis*-targeting phages have genomes that are 30-150kb in length and belong to the class Caudoviricetes, which encompasses tailed, double-stranded DNA viruses^26^. The phages we isolated were also predicted to belong to Caudoviricetes and appear to belong to four different genera, only one of which, *Efquatrovirus*, has been previously described. *Efquatrovirus* phages have been primarily isolated from wastewater and sewage^27^, and they have emerged as leading candidates for enterococcal phage therapy due to their high specificity and efficacy against antibiotic-resistant *Enterococcus* strains. Though some phages exhibiting cross-species infectivity within the genus *Enterococcus* have been described^20,29^, the host range of most *E. faecalis* phages is narrow and often limited to specific strains. The phages we isolated showed varied infectivity profiles against the tested EFIE clinical isolates, with the most active phage only able to infect 8/18 (44%) isolates. This strain-restricted activity, coupled with the fact that none of the isolated phages could infect any of the *E. faecium* isolates we tested, suggest that these phages are likely specific for *E. faecalis*.

Combining phages with complementary activities together into cocktails is a promising approach to mitigate the narrow host range of many phages^17^, and several studies have reported that both β-lactams and daptomycin can enhance phage efficacy against enterococci^21,22,28^. Similarly, we found that a three-phage cocktail consisting of EFS07, EFS13, and EFS15 was active against nearly 90% of EFIE clinical isolates tested, and there was no apparent antagonism between the phages when they were combined together. Additionally, when we combined both individual phages and the three-phage cocktail with subinhibitory concentrations of ampicillin or daptomycin, we observed enhanced bacterial inhibition and killing. A prior study defined phage-antibiotic synergy as at least a 100-fold reduction in bacterial abundance in the combination group compared to the most effective monotherapy, while a reduction between zero and 100 is considered an additive effect^30^. We found that most phage-antibiotic combinations at the concentrations we tested exhibited additivity when assessed after 20 hours, although the extent of bacterial inhibition varied depending on the antibiotic used and the *E. faecalis* isolate tested. Ampicillin inhibits cell wall synthesis by targeting penicillin-binding proteins (PBPs) to weaken the peptidoglycan layer, which may enhance phage adsorption by exposing or modifying cell surface receptors and facilitating phage DNA injection due to compromised cell wall integrity^31^. Daptomycin disrupts cell membranes with the assistance of calcium^32^, though the mechanism by which it might improve phage activity is unclear. Nonetheless, our findings suggest that the phages we isolated could be suitable partner agents alongside the antibiotics currently used to treat EFIE.

Even though phage therapy is being increasingly used in humans and phages appear to have a favorable safety profile, testing of phage efficacy using relevant *in vivo* models is still an important step in preclinical development. We therefore tested the efficacy of our three-phage cocktail in *G. mellonella*, an invertebrate model for studying microbial pathogenesis and antimicrobial efficacy, including infections caused by *E. faecalis*^33^ and treatment with phage therapy^34^. *G. mellonella* possess an innate immune system with structural and functional parallels to mammals, including hemocytes (analogous to neutrophils) and the production of antimicrobial peptides^35^. We selected a highly virulent ST6 isolate for infection and found that treatment with the phage cocktail two hours after infection significantly improved larval health and reduced mortality, though the effect was partial and did not restore survival to the level of the injection control condition. There could be several reasons for this, including poor phage distribution, phage inactivation by host proteases, irreversible damage to host tissue prior to phage administration, or the development of phage resistance. Although full recovery in the phage-treated larvae was not achieved, our results nonetheless demonstrate the therapeutic potential of our three-phage cocktail for treating EFIE *in vivo*.

This study had several limitations. First, we tested a limited number of phages and only screened EFIE clinical isolates from a single center, raising the possibility that our results might not be generalizable to other settings. Additionally, while we assessed phage-antibiotic combinations *in vitro*, we treated infected *G. mellonella* larvae with phage alone, which prevented us from being able to assess the *in vivo* efficacy of combination therapy in this model. Additionally, *G. mellonella* lack adaptive immunity, and phage pharmacokinetics and pharmacodynamics in this model might differ from those in vertebrates^35^. Finally, we observed different levels of phage susceptibility in the DVT1549 isolate depending on the assay used, highlighting disagreements between *in vitro* susceptibility testing methods and a need for more studies comparing *in vitro* assessments with *in vivo* phage activity. Future studies could also employ vertebrate models of infection that are more translationally relevant to more accurately assess the *in vivo* efficacy of *E. faecalis*-targeting phage therapy for further preclinical development.

In summary, we characterized 10 novel *E. faecalis*-targeting phages from municipal wastewater, found that a three-phage cocktail in combination with either ampicillin or daptomycin has enhanced activity against EFIE clinical isolates, and showed that phage treatment can improve survival of a lethal infection in *G. mellonella*. These findings demonstrate the potential of phage-based therapies as effective adjuncts for treating *E. faecalis* infections.

## Materials and Methods

### Bacterial and phage isolation

*E. faecalis* clinical isolates were collected from blood cultures of patients with infective endocarditis (IE) as identified per modified Duke’s criteria^36^ at University of Pittsburgh Medical Center (UPMC)-affiliated hospitals in Pittsburgh, Pennsylvania from 2018 to 2023, and were cryopreserved at –80°C in Brain Heart Infusion (BHI, BD Bacto™) with 16% glycerol. Collection of EFIE clinical isolates was approved by the Institutional Review Board at the University of Pittsburgh School of Medicine (STUDY22050046). Bacteria were grown in Todd Hewitt Broth (THB) or Mueller Hinton Broth (MHB) supplemented with 10 mM MgSO_4_ unless stated otherwise. For experiments with daptomycin, cation-adjusted MHB containing 50 μg/mL CaCl_2_ was used. Phages were isolated from untreated municipal wastewater collected from the greater Pittsburgh area. Briefly, an aliquot of an overnight bacterial culture of an *E. faecalis* clinical isolate grown in magnesium-supplemented THB was pelleted and resuspended in sterile SM buffer (50 mM Tris-HCl, 100 mM NaCl, 8 mM MgSO; pH 7.5). Concentrated, sterile-filtered wastewater was added to the resuspended culture and incubated at room temperature for at least 15 minutes. The mixture was then added to molten magnesium-supplemented THB top agar (0.35% agar supplemented with 10 mM MgSO_4_) and spread over a pre-warmed THB bottom agar (1.5% agar supplemented with 10 mM MgSO_4_) plate. Once solidified, plates were incubated overnight at 37°C. The following day, plaques were picked with a sterile pipet tip, and the phage was eluted into a small volume of SM buffer overnight at 37°C. Eluted phage was then spot-titered back onto a lawn of the original *E. faecalis* isolate. The following day a plaque with consistent morphology was again picked and eluted. Each phage underwent at least three rounds of plaque purification to ensure a monophage preparation. Isolated phages were stored in SM buffer at 4°C.

### Whole genome sequencing

Genomic DNA was previously extracted from each isolate using a DNeasy Blood & Tissue Kit (QIAGEN, Germantown, MD, USA) and sequenced on the Illumina platform using 150-bp paired-end reads^10^. *E. faecalis* genomes were assembled de novo using CLC Genomics Workbench v23.0.5, annotated with Prokka v1.14.5^37^ and compared to each other with Roary v3.11.2^38^. A midpoint-rooted phylogenetic tree of EFIE isolates was constructed from the single-copy core genome alignment using RAxML v8 with the GTRCAT algorithm and 100 iterations^39^. Multi-locus sequence types (STs) were identified with PubMLST^40^, and single locus variants (SLVs) were grouped with the closest ST. Genomic DNA was extracted from 200 μL of phage lysate also using the DNeasy Blood & Tissue Kit (QIAGEN, Germantown, MD, USA). If the extracted DNA content was low, multiple 200 μL extraction reactions were run over a single DNA-binding column to increase yield. Genomes were sequenced on the Illumina platform using 150-bp paired-end reads and were assembled using SPAdes v3.14.1 with the –metaviral flag^41^. A viral dendrogram was constructed using ViPTree v4.0^42^, and phage lifestyle was predicted by PhageAI^24^.

### Generation of high-titer phage lysates

Phages were propagated using the original *E. faecalis* host used for phage isolation. Overnight cultures grown in THB at 37 °C were centrifuged at 13,000 × g for 1 min and pellets were resuspended in SM buffer. Phages were mixed with 100 µL of host cells in SM buffer with different volumes of phage (10, 25, or 50 µL), incubated at room temperature for 15 min, then combined with 10 mL of molten THB top agar and poured onto THB bottom agar plates. Plates were incubated overnight at 37 °C. Plates showing webbed lysis were flooded with 10 mL SM buffer and incubated for 1 h. The lysate was centrifuged at 4,000 × g for 15 min, and supernatants were filtered (0.22 µm) to obtain sterile, high-titer phage stocks.

### Phage titering

Phage titers were determined using a double-layer overlay assay^43^. Briefly, bacterial lawns were prepared by mixing 5 mL of molten THB top agar and 100 µL of resuspended host culture in SM buffer, then poured onto square THB bottom agar plates. Phage lysates were 10-fold serially diluted in SM buffer and 5 µL of each dilution was spotted onto the solidified lawns followed by incubation overnight at 37°C. Plaques were counted at dilutions yielding 10–100 plaques, and titers were calculated as: PFU/mL = #plaques × 200 × dilution factor. Efficiency of plating (EOP) was used to compare phage infectivity across hosts: EOP = [Titer on clinical isolate] / [Titer on host]. An EOP ∼1 indicated similar infectivity as the host isolate, whereas lower values indicated reduced activity on the clinical isolate.

### Modified phage-antibiotic synergy (PAS) assay

A modified PAS assay was adapted from previous studies^30^. Phage stocks were prepared at 1×10 PFU/mL. Log-phase bacterial cultures were diluted to OD = 0.003 in 5 mL MHB (∼10 CFU/mL). Four treatment groups were tested for each isolate: antibiotic only, phage only, antibiotic-phage combination, and untreated. Ampicillin and daptomycin were used at 0.5 µg/mL, which was ½ the MIC for most isolates. Phages were added at MOI = 10 (∼10 PFU/mL) for host isolates and at MOI = 100 (∼10^8^ PFU/mL) for EFIE clinical isolates due to the low EOP of the latter. At 0 h, 100 µL from the untreated group was serially diluted 10-fold from 10^0^ to 10 and 10 µL of each dilution was spotted on MHB agar. At 24 h, 0.5 mL samples from all groups were collected, washed with PBS, and resuspended. Then 100 µL from each sample was serially diluted to 10 and plated. Plates were incubated overnight at 37 °C, and colonies were counted to determine CFU. CFU ratio = CFU / CFU. For growth curves, 200 µL of each treatment at 0h was added to 96-well plates (8 replicates). Plates were sealed and incubated at 37 °C in a Synergy H1 plate reader with OD measured every 30 min for 20 h, including 2.5 min of shaking at 200 rpm before each reading. Background-subtracted OD values were recorded, with negative values recorded as 0.

### In vivo Galleria mellonella larvae model

*Galleria mellonella* larvae were purchased from Grubco (Fairfield, OH, USA) and used as an *in vivo* infection model as described previously^33^. Healthy larvae (0.22–0.32 g) without signs of pupation, discoloration, or desiccation were selected, randomly assigned to experimental groups, and stored at 4 °C for up to 48 h before use. For preliminary screening of *E. faecalis* isolates, experimental groups (n = 20 larvae per group) consisted of a PBS-injected control group and infection groups challenged with isolates DVT958, DVT1548, DVT1549, and DVT1720. Bacterial cultures were grown overnight in THB at 37 °C, diluted 1:100, incubated for 3 h, adjusted to OD = 0.2, and further diluted 1:100 in PBS (∼10 CFU/mL). Final concentrations were verified by plating and CFU enumeration. Larvae were chilled for 30 min before injection with 5 µL of bacterial suspension or PBS into a proleg. Following injection, larvae were incubated at 37 °C, and survival was monitored every 24 h for three days. Health status was further evaluated using the Pathogenicity Score Index (PSI)^25^, which assesses movement and melanization, with higher scores indicating poorer health. For further experiments using isolate DVT1549 only, groups (n = 30 larvae per group) included PBS control, bacterial infection, and phage-treated infection. Procedures were identical to those described above, except that a higher bacterial burden (5 μL of ∼2×10 CFU/mL) was used to increase the lethality, and at 2 hours post-infection larvae were injected with either 5 µL PBS for the PBS control and bacterial infection groups) 5 µL of phage cocktail (∼2×10 PFU/mL, MOI = 100).

## Statistical analysis

Differences in AUC and log_10_[CFU ratio] of PAS testing and PSI were assessed via one-way ANOVA followed by Tukey’s multiple comparisons test. Kaplan-Meier survival curves of *Galleria mellonella* larvae among different groups were compared via log-rank tests.

## Data Availability

Whole genome sequencing data of phages, *E. faecalis* and *E. faecium* isolates were deposited in NCBI with accession numbers listed in Table 1 and Table S1.

## Acknowledgements

We gratefully acknowledge all members of the Van Tyne lab for their helpful input throughout the conduct of this study and preparation of the manuscript. We also acknowledge Dr. Gina Suh at the Mayo Clinic for providing clinical *E. faecalis* isolates. This work was supported by the Department of Medicine at the University of Pittsburgh School of Medicine. Y. L. was also supported by the Pitt-Tsinghua Partnership Program. The funders had no role in study design, data collection and analysis, preparation of the manuscript, or decision to submit the manuscript for publication.

## Figure Legends

**Figure S1.** Quantitative CFU analysis of host isolates after 24 hours of single agent or combination therapy. Three *E. faecalis* host isolates were treated with (A) ampicillin (AMP) 0.5 μg/mL, phage at MOI 10, and combination therapy, or (B) daptomycin (DAP) 0.5 μg/mL, phage at MOI 10, and combination therapy, with growth control included. CFU ratio = CFU_24h_/CFU_0h_. Mean values were shown as bars. Statistical significance was determined by one-way ANOVA followed by Tukey’s multiple comparisons test. *p< 0.05, **p<0.01, ***p< 0.001, ****p<0.0001.

