## Supplemental Fig. 1 for "Evaluation of phage activity against diverse *Enterococcus faecalis* from infective endocarditis"

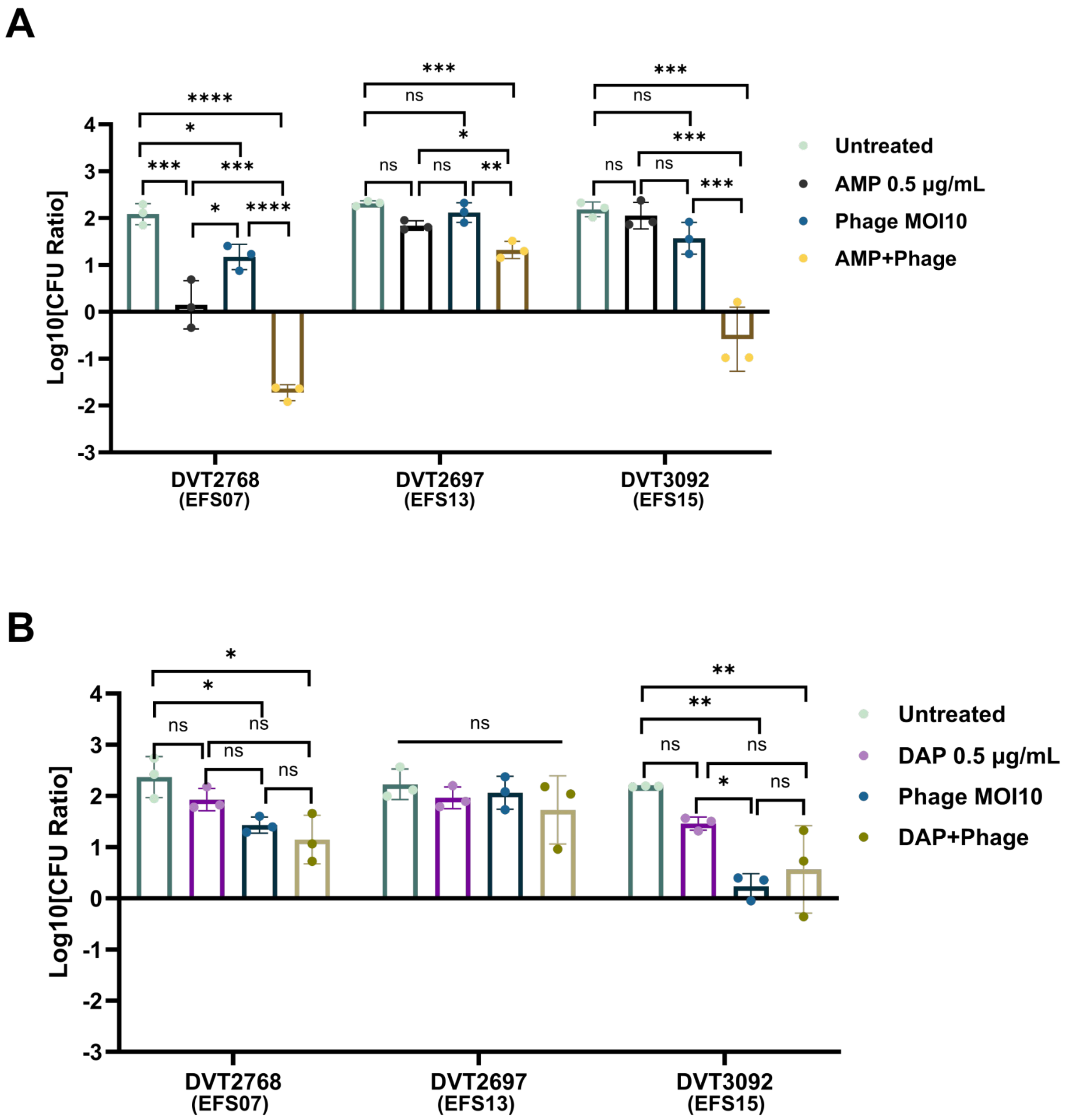


**Figure S1. Quantitative CFU analysis of host isolates after 24 hours of single agent or combination therapy.** Three *E. faecalis* host isolates were treated with (A) ampicillin (AMP) 0.5 µg/mL, phage at MOI 10, and combination therapy, or (B) daptomycin (DAP) 0.5 µg/mL, phage at MOI 10, and combination therapy, with growth control included. CFU ratio = $CFU_{24h}/CFU_{0h}$. Mean values were shown as bars. Statistical significance was determined by one-way ANOVA followed by Tukey's multiple comparisons test. $^{*}p < 0.05$, $^{**}p < 0.01$, $^{***}p < 0.001$, $^{****}p < 0.0001$.
